# AutoGaitA: A versatile quantitative framework for kinematic analyses across species, perturbations and behaviours

**DOI:** 10.1101/2024.04.14.589409

**Authors:** Mahan Hosseini, Ines Klein, Veronika Wunderle, Carolin Semmler, Taylan D. Kuzu, Ann-Kathrin Kramer, Marianna Tolve, Vlad Mardare, Ana Galvao, Moritz Haustein, Christian Grefkes, Tatiana Korotkova, Ansgar Büschges, Gereon R. Fink, Peter H. Weiss, Silvia Daun, Graziana Gatto

## Abstract

Individual behaviours require the nervous system to execute specialised motor programs, each characterised by unique patterns of coordinated movements across body parts. Deep learning approaches for body-posture tracking have facilitated the analysis of such motor programs. However, translating the resulting time-stamped coordinate datasets into meaningful kinematic representations of motor programs remains a long-standing challenge. We developed the versatile quantitative framework AutoGaitA (Automated Gait Analysis), a Python toolbox that enables comparisons of motor programs at multiple levels of granularity and across tracking methods, species and behaviours. AutoGaitA allowed us to demonstrate that flies, mice, and humans, despite divergent biomechanics, converge on the age-dependent loss of propulsive strength, and that, in mice, locomotor programs adapt as an integrated function of both age and task difficulty. AutoGaitA represents a truly universal framework for robust analyses of motor programs and changes thereof in health and disease, and across species and behaviours.

## Introduction

Animals and humans employ an impressively rich array of motor behaviours, such as grooming, swimming or walking, each requiring coordinated neural activity across the nervous system to execute the appropriate motor program^1,2^. Individual motor programs are executed with redundant kinematic solutions, which arise from distinct combinations of joint positions, angles and velocities^3^ and operate within species-specific biomechanical constraints^1,2^. Consequently, the kinematic outputs of even seemingly simple motor programs, such as walking, exhibit tremendous variability, which complicates deciphering the underlying neural code.

Advances in deep-learning methods, such as DeepLabCut (DLC^4^) and SLEAP^5^, have considerably improved our ability to track body landmarks across behavioural tasks and species. Nevertheless, no universal framework currently exists to coalesce these time-stamped body coordinates into meaningful representations of motor programs. Current analysis methods rely on commercial software (e.g., DigiGait, Motorater, Theia3D, SIMI) or custom scripts that are often task- and/or species-specific, limiting their scope. Toolboxes assessing rodent kinematics are task-specific, being constrained to ladder, treadmill (ALMA^6^; PMotion^7^) or beam walking (Forestwalk^8^; Ledged Beam Walking^9^). Further, available analyses provide only basic gait parameters (PMotion, Forestwalk), disease-specific features, e.g. for stroke (PMotion), or complex kinematic extrapolations (ALMA, Ledged Beam Walking). Additionally, although numerous species-specific approaches to gait analysis have been developed for fruit flies^10–13^, mice^6,8,9,14,15^ and humans^16^, no framework exists for cross-species kinematic comparisons.

AutoGaitA (Automated Gait Analysis) provides a versatile framework to standardise the analysis of body coordinates and assess motor programs by comparing kinematic features at different levels of granularity, from individual landmarks to full-body coordination strategies. Using AutoGaitA, we here demonstrate that fruit flies, mice and humans, despite their divergent biomechanics, share a common age-dependent adaptation strategy: they reduce leg propulsive strength during walking to gain postural stability. Furthermore, we showed that mice employ distinct strategies, defined by age and task difficulty, to preserve the flexibility and robustness of motor execution, suggesting that adaptation strategies emerge as integrative functions of concomitant perturbations.

In sum, AutoGaitA fills a critical gap in the methods currently used to study motor control policies by providing a general-purpose toolbox to assess and compare motor programs and their changes in health and disease across species and behaviours.

## Results

### A universal framework to analyse rhythmic motor programs at different granularity levels

The heterogeneity of behaviours, model organisms and tracking methods has so far hindered the development of a standardised quantitative framework for the kinematic analysis of motor programs. AutoGaitA addresses this methodological gap by providing a universal framework applicable to any rhythmic behaviour and species. We developed three first-level toolboxes, implementing the same overall workflow to accommodate the diversity of tracking methods (Fig. 1a-c). AutoGaitA DeepLabCut (DLC) and AutoGaitA SLEAP analyse 2D coordinates obtained with DLC^4^ or SLEAP^5^, respectively. AutoGaitA Universal 3D analyses 3D coordinates tracked with any marker-based or marker-less method.

**Figure 1.**
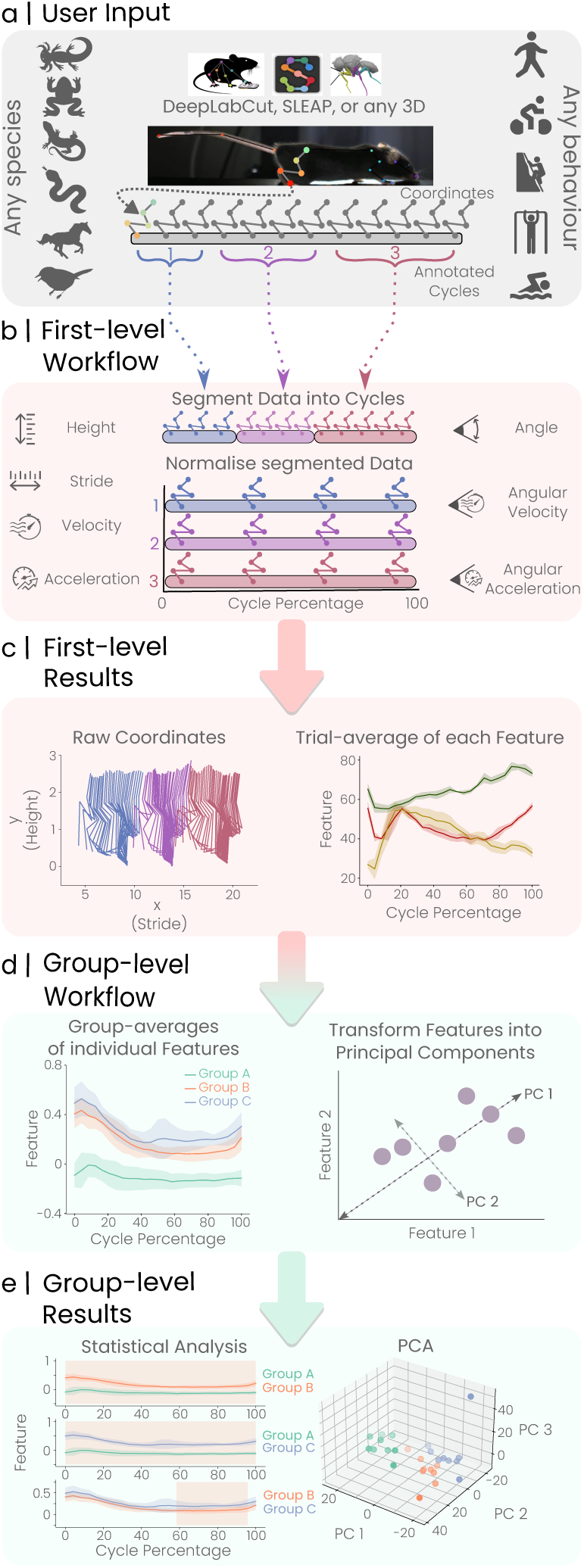
AutoGaitA’s universal framework for the kinema9c comparisons of rhythmic behaviours within and across species. **a**, User input. AutoGaitA loads 2D (from DeepLabCut or SLEAP tracking) or 3D (any 3D method) body coordinate data as well as the annotation table storing Fme information. The AutoGaitA workflow is applicable to any model organisms (e.g. axolotl, salamander, frogs, songbirds) performing a variety of rhythmic behaviours. **b**, First-level workflow. Landmark coordinates are first segmented into cycles based on the annotation table and then normalised. Normalised kinematic features (stride, height, angles, velocities and accelerations) are computed and averaged for each trial. **c**, First-level results. Results are generated as image and tabular files, summarising raw coordinates and kinematic features for each trial aPer segmentation, normalisation, and averaging. **d**, Group-level workflow. Comparisons of kinematic features among groups are computed via statistical tests and principal component analysis (PCA). **e**, Group-level results. Results of statistical tests (e.g., pair-wise Fme points of statistical significance and p-values) are provided for each variable tested. PCA results are provided as customisable scaQerplots, videos, and tabular files.

AutoGaitA’s first-level toolboxes transform time series of body coordinates into standardised sets of kinematic outputs. First, AutoGaitA loads the coordinate tables containing the time series of landmark positions and the manually generated annotation tables specifying the timestamps of all behaviour-cycles of interest (Fig. 1a). In this context, a behaviour-cycle (henceforth cycle) defines the simplest instance of a rhythmic behaviour with clearly defined start and end timepoints. For example, a step is the cycle of walking, starting when the foot is lifted off the ground and ending just before the foot is lifted again for the next step. The coordinates are segmented into user-annotated cycles (Fig. 1b), and the segmented data is processed to compute kinematic features: stride (horizontal distance), height (vertical distance), angles, velocities, and accelerations of any user-defined landmark. Next, the segmented data is normalised to bring individual cycles, naturally varying in duration, to a fixed length, making the cycles’ kinematics well-suited for comparing and averaging. The generated first-level results consist of image and tabular files containing raw, segmented, normalised and averaged data (Fig. 1c).

The AutoGaitA Group toolbox computes and compares group-level effects after averaging the first-level results of each group (Fig. 1d). Individual kinematic features can be assessed statistically with a permutation test, an ANOVA, and Tukey’s test, while more general kinematic patterns can be identified using principal component analyses (PCA). Datasets for each group are provided as tabular files, enabling custom post-hoc analyses when required. The results of group-level comparisons are provided as image and tabular files (Fig. 1e).

To demonstrate AutoGaitA’s versatility and applicability, we analysed a published dataset of tracked body landmarks in human subjects performing distinct rhythmic movements (MoVi dataset^17^). Using AutoGaitA, we were not only able to analyse a range of rhythmic behaviours - from running to jumping - but also to cluster the underlying motor programs in PCA space, with similar movements such as walking and running being in closer proximity (Supplementary Note 1, Supplementary Fig. 1).

### Species-specific and convergent strategies of age-dependent motor adaptation

Comparing motor programs across species has long posed a significant challenge due to fundamental biological differences in morphology, biomechanics and size, as well as the heterogeneity of experimental paradigms and posture tracking methods. Leveraging AutoGaitA’s ability to analyse coordinate data independently of the model organism and tracking algorithm, we assessed the motor programs underlying walking (henceforth locomotor program) - one of the most fundamental rhythmic behaviours - in flies (*Drosophila Melanogaster*), mice (*Mus Musculus*) and humans. DLC was used to track body landmarks in flies walking on a spherical treadmill, mice walking on a 25-mm wide beam, and humans walking on a wide walkway (Fig. 2a-c). We focused our analyses on posterior/lower limb landmarks: the thorax-coxa (Th-Cx), the coxa-trochanter (Cx-Tr), the femur-tibia (Fe-Ti), the tibia-tarsus (Ti-Tar), and the tip of the tarsus (Tar) on the fly posterior leg (Fig. 2d); the iliac crest, hip, knee, ankle and middle hindpaw on the mouse hindlimb (Fig. 2e); and the hip, knee, ankle, and midfoot on the human leg (Fig. 2f). We analysed how the kinematic features of these landmarks vary during the step cycle, defined from the beginning of swing (foot is lifted from the ground) to the end of stance (just before the foot is lifted from the ground for the next step) (Fig. 2f). The stick diagrams, illustrating the horizontal (stride) and vertical (height) displacement of the limbs during a step cycle, revealed the divergent kinematics resulting from species-specific biomechanics (Fig. 2g-i).

**Figure 2.**
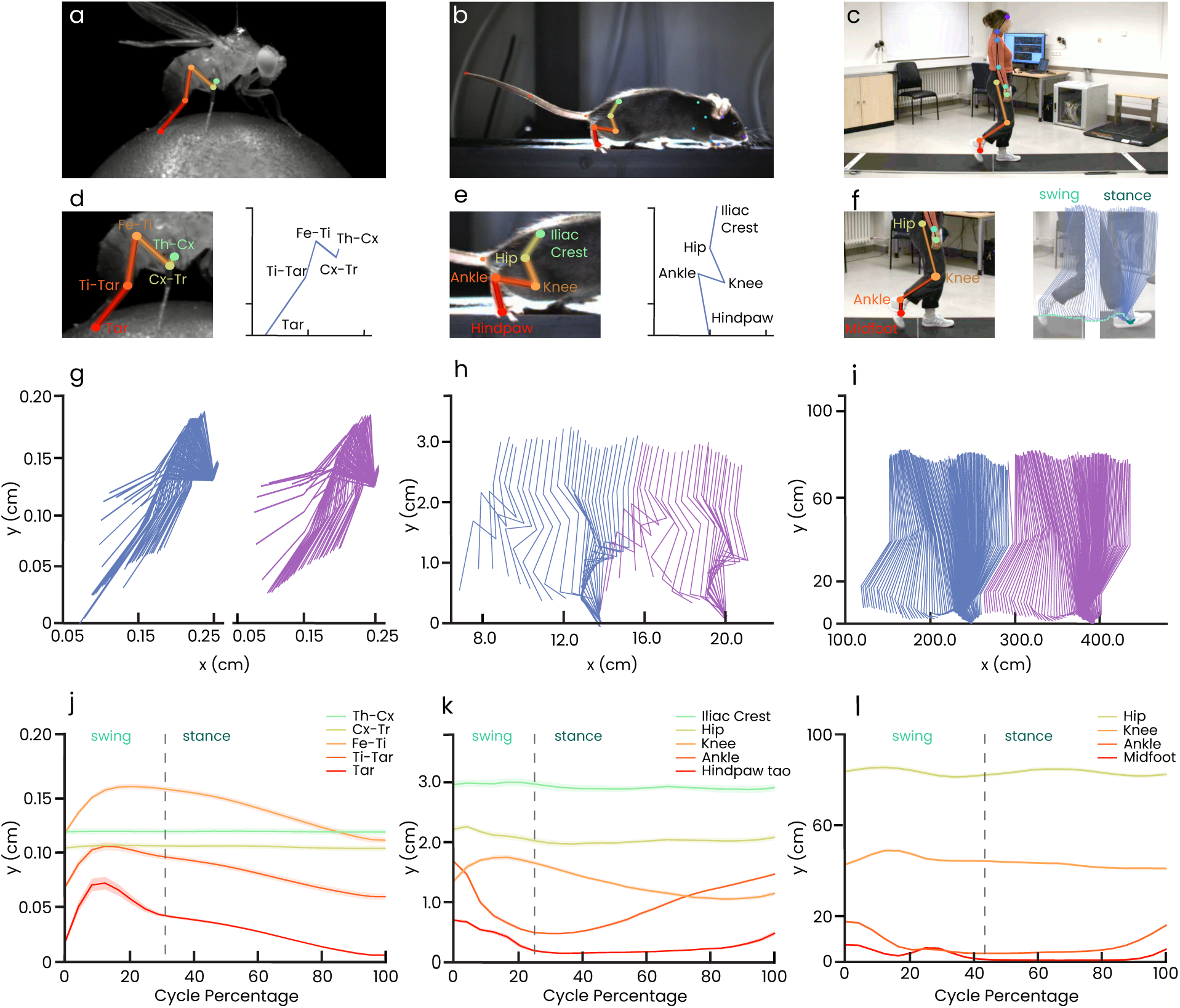
Divergent and convergent kinema9c strategies underlying locomotor programs in flies, mice, and humans. **a-c,** Snapshots of a fly walking on a spherical treadmill (a), a mouse walking on a 25 mm wide beam (b), and a human walking on a walkway (c). **d-f,** Snapshots illustrating the landmarks analysed on the fly posterior leg (d), the mouse hindlimb (e) and the human leg (f). The right panels in d-e illustrate how landmarks translate to the stick diagrams below. The right panel in f depicts a step cycle and its division into swing (toe-off) and stance (toe-down) phases. **g**, Stick diagram illustrating the horizontal and vertical displacements of the fly posterior leg. Note that flies were tethered. **h-i**, Stick diagrams of the mouse hindlimb (h) and human leg (i) for two consecutive steps. **j**, Group-averaged variations in the height of key landmarks of the fly posterior leg throughout the step-cycle showing that all distal joints are moved almost synchronously upward and downward to generate propulsion. **k**, Group-averaged variations in the height of key landmarks of the mouse hindlimb throughout the step-cycle showing that the hip provides the initial power stroke, while the ankle and knee invert their relative heights to generate the subsequent thrust. **l**, Group-averaged variations in the height of key landmarks of the human leg throughout the step-cycle showing that in humans the hip moves pendulously to ensure stability, while distal joints, particularly the ankle, generate propulsion. Data are presented as mean±SEM, SEM is shown as shaded areas. N=6 for flies, N=9 for mice, N=29 for humans.

In tethered flies, the proximal Th-Cx and Cx-Tr remained in relatively fixed positions, while the distal Fe-Ti, Ti-Tar and Tar moved, almost synchronously, upward during swing and downward during stance (Fig. 2g,j, Supplementary Fig. 2d). This resulted in all distal joints being maximally flexed at the swing-to-stance transition and maximally extended at the stance-to-swing transition (Supplementary Fig. 2a), suggesting that in flies, distal joints are synergistically coordinated to provide the primary power stroke for forward propulsion of the posterior leg. In mice, the iliac crest remained relatively static throughout the step cycle, the hip moved upward and downward only during swing, the knee and the hindpaw moved upward during swing and at the end of stance and downward during mid-stance, while the ankle exhibited opposite behaviour, moving downward during swing and upward during stance (Fig. 2h,k, Supplementary Fig. 2e). This resulted in the ankle, knee and hip being similarly maximally flexed mid-swing, but diverging in their peak extension, with the ankle and knee being maximally extended at the beginning of stance and the hip at the end of stance (Supplementary Fig. 2b). Together, this kinematic pattern suggests that in mice, the hip provides the initial power stroke, while the relative inversion of ankle and knee positions drives the forward thrust. In humans, the hip oscillated through the step cycle, marking the transitions across swing and stance, the knee moved upward during swing and downward during stance, while the ankle and the midfoot oscillated during swing, and moved upward at the end of stance (Fig. 2i,l, Supplementary Fig. 2f). This resulted in the peak flexion and extension of the knee occurring during swing and of the ankle at the end of stance and beginning of swing (Supplementary Fig. 2c). Taken together, this pattern suggests that in humans, the ankle push-off at the end of stance generates the additional power stroke that, together with the hip-driven oscillations, promotes the forward propulsion. Notably, our kinematic analysis suggests that the generation of propulsive strength relies on coordinated flexion/extension of all distal joints in flies, whereas it requires opposite displacement of knee, ankle and hindpaw in mice, and hip oscillations and ankle push-off in humans. Despite distinct biomechanics, all species converge on landmark velocities increasing along the proximal-to-distal axis (Supplementary Fig. 2g-i). Thus, leveraging the numerous kinematic features assessed by AutoGaitA, we revealed that the mechanisms underlying locomotor programs in hexapedal (flies), quadrupedal (mice) and bipedal (human) model organisms converge in setting a proximal-to-distal gradient of landmark velocities and diverge in how leg propulsive strength is generated.

Next, we investigated how the same physiological perturbation, ageing, affects the execution of locomotor programs across species. The PCA obtained with the AutoGaitA group-level analysis indicates that the locomotor program shows an ageing signature in each species (Fig. 3a-c). Remarkably, not all older subjects segregate from the “young cluster”, showing inter-individual variability in response to ageing in all species (Fig. 3a-c). To assess the components driving the age-dependent adaptation of the locomotor programs, we analysed individual features next. Intriguingly, and consistent with their species-specific roles in generating propulsion, we observed that age reduced i) in flies, the flexion of the Fe-Ti and Cx-Tr angles at the swing-stance transition, shortening the power stroke length (Fig. 3d, Supplementary Fig. 3a); ii) in mice, the extension of the ankle at the swing-stance transition, affecting the knee/ankle dynamics driving the forward thrust (Fig. 3e, Supplementary Fig. 3b); iii) in humans, the ankle flexion during stance that usually provides the additional push-off power to move forward (Fig. 3f). Taken together, these observations revealed a conserved effect of aging on locomotor programs: a reduction in the generation of propulsive strength resulting from changes in the species-specific kinematic features that underlie the forward thrust of the leg.

**Figure 3.**
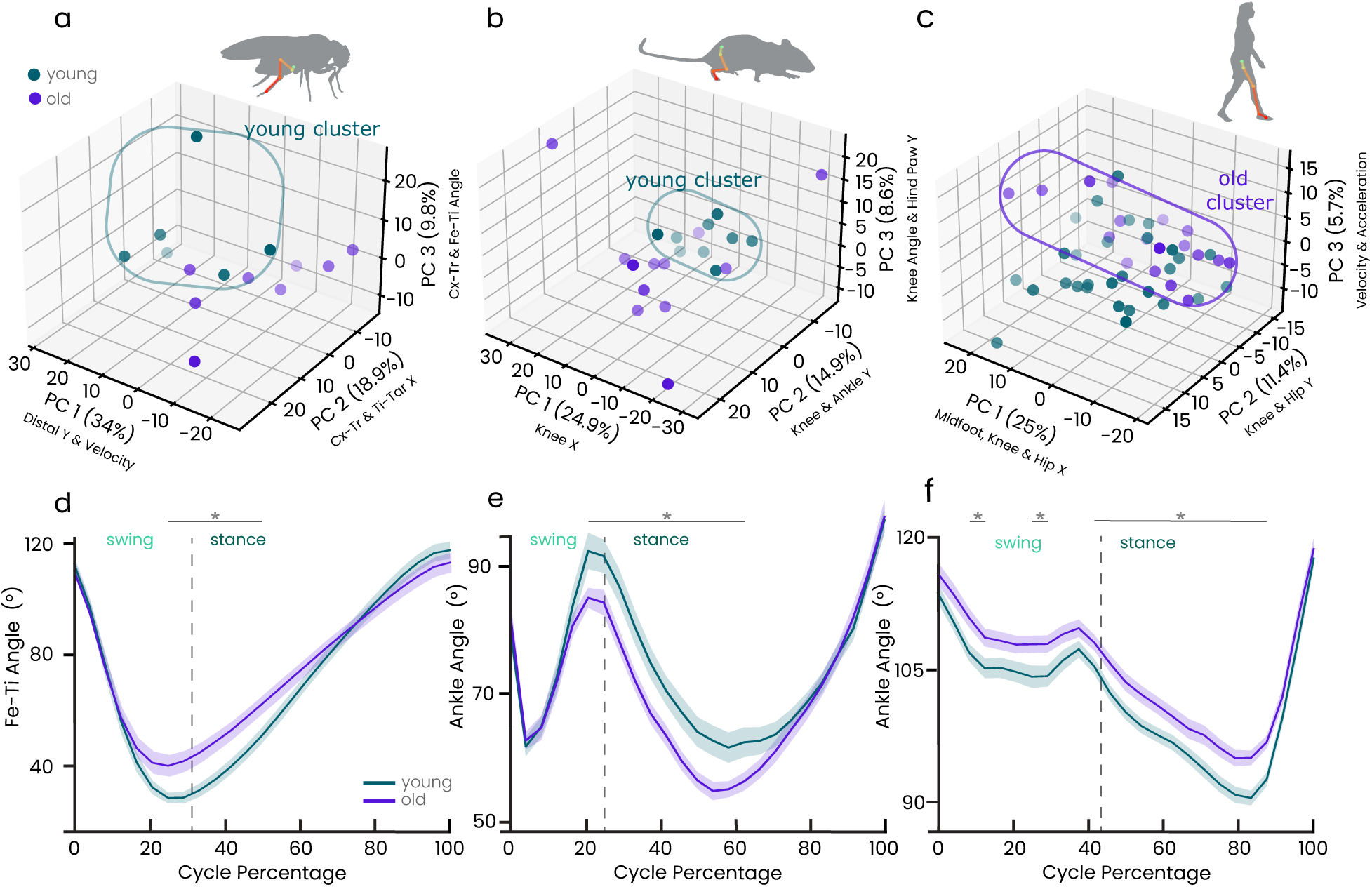
Age-dependent adapta9on of locomotor programs across species. **a**, PCA scaQerplot of fly posterior leg kinematics shows segregated clusters for young (green, 2-3 days, N=6) and old (purple, 21-22 days, N=6) flies. Each circle represents an individual fly. **b**, PCA scaQerplot of mouse hindlimb kinematics shows segregated clusters for young (8 months, N=9) and old (24 months, N=12) mice. Each circle represents an individual mouse. **c**, PCA scaQerplot of human leg kinematics shows partially segregated clusters for young (21-36 years, N=29) and old (46-85 years, N=18) humans. Each circle represents a single participant. **d**, Flexion of the femur-tibia (Fe-Ti) angle at the swing-stance transition is reduced in old compared to young flies. **e**, Extension of the ankle angle at the swing-stance transition is reduced in old compared to young mice. **f**, Flexion of the ankle angle during stance is reduced in old compared to young humans. Data are presented as mean±SEM, SEM is represented as shaded areas. Statistical analysis: one-way ANOVA followed by Tukey’s post-hoc test. Significance is indicated with asterisks, *p<0.05. See Supplementary Tables S1-S9 for complete statistical and PCA results.

### AutoGaitA accurately detects task- and age-dependent changes in mouse kinematics

Real-world locomotion rarely involves a single, discrete perturbation, but is rather influenced by multiple, simultaneous disturbances, whose effects add up to more than their sum. Thus, the nervous system must deploy adaptive motor programs that account for interactions between different perturbations. We therefore investigated how the adaptation of locomotor programs to increased task difficulty evolves with age, studying how untrained 3-, 8-, and 24-month-old C57BL6/J mice cross beams of varying widths: 25-, 12-, and 5-mm (Fig. 4a-4c).

**Figure 4.**
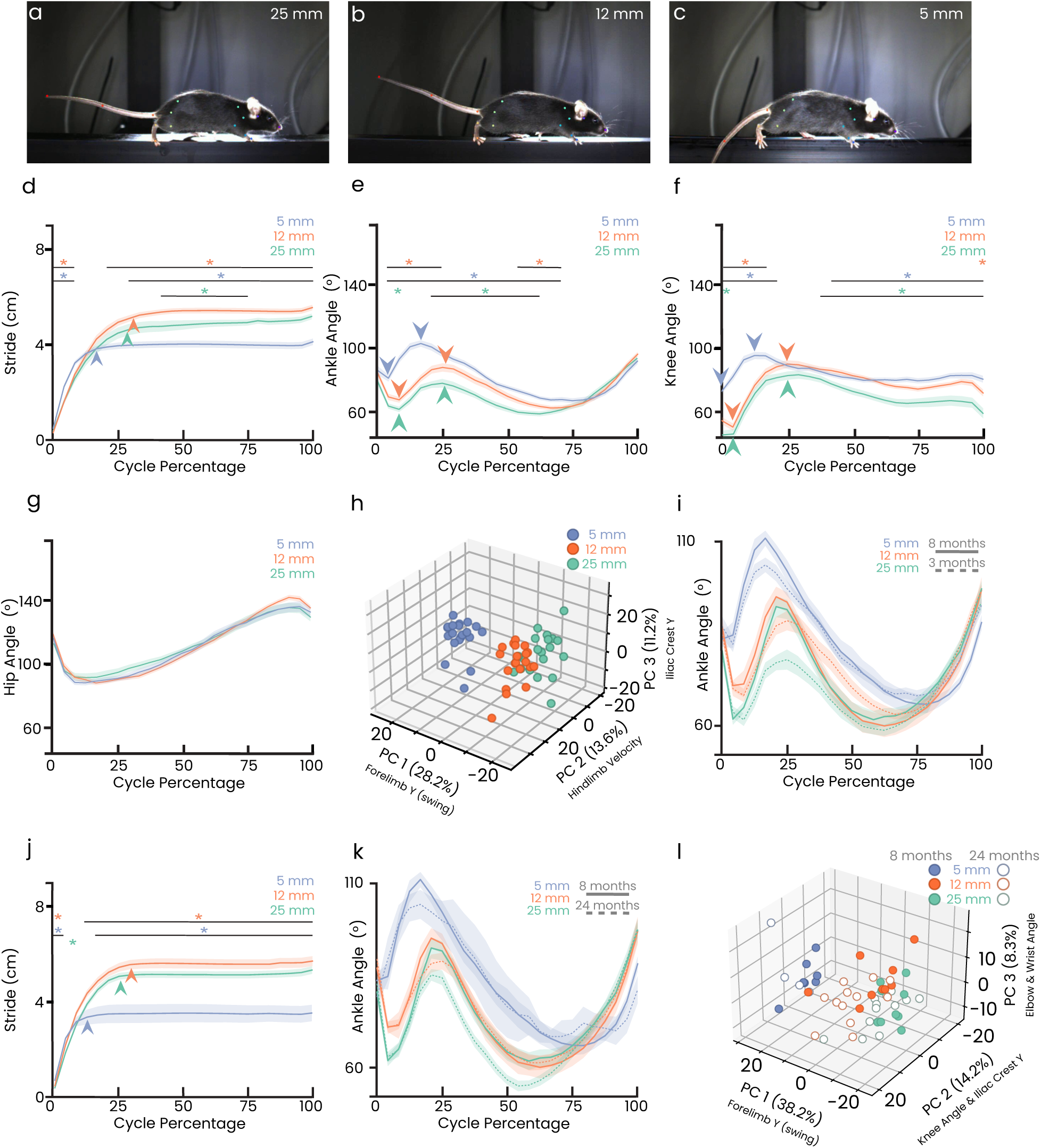
Task- and age-dependent motor adapta9on of locomotor programs in mice. **a-c**, Snapshots of a mouse crossing the 25 mm (a), 12 mm (b) and 5 mm (c) wide beam. **d**, 3-month-old mice shorten their stride when crossing a 5 mm wide beam (blue) compared to the 25 mm (green) and 12 mm (orange) wide beams. The shortened stride also corresponds to a shorter swing, as indicated by the arrowheads coloured according to beam widths. **e**-**g** Changes during the step cycle in the ankle (e), knee (f) and hip (g) angles in 3-month-old mice crossing the 25 mm (green), 12 mm (orange) and 5 mm (blue) wide beams. Mice crossing the narrowest (5 mm) beam showed reduced flexion and increased extension of the ankle and knee during swing, as indicated by the coloured arrowheads. **h**, PCA of changes in heights, angles and velocities of the forelimb and hindlimb throughout the step cycle revealed segregated clusters based on beam widths. Each circle represents an individual mouse. **i**, Ankle angles of 8-month-old (solid lines) and 3-month-old (dashed lines) mice crossing the three beams. 8-month-old mice exhibited a similar kinematic paQern of beam-induced adaptation, but an even more accentuated extended-limb posture. **j**, 24-month-old mice shorten their stride even more when crossing a 5 mm wide beam (blue) compared to the 25 mm (green), 12 mm (orange) wide beams. The shortened stride also corresponds to a shorter swing, as indicated by the arrowheads coloured according to beam widths. **k**, Ankle angles of 8-month-old (solid lines) and 24-month-old (dashed lines) mice crossing the three beams. 24-month-old mice exhibited a similar kinematic paQern of beam-induced adaptation, but flexed their joints more, indicating a crouched position on the beam to lower their centre of mass. **l**, PCA of changes in heights, angles and velocities of the forelimb and hindlimb throughout the step cycle between 8-month-old (solid lines) and 24-month-old (dashed lines) mice revealed segregated clusters based on beam widths and age. Each circle represents an individual mouse. Data presented as mean±SEM. N=20 for 3-month-old, N=9 for 8-month-old and N=12 for 24-month-old mice. Statistical comparison: one-way ANOVA followed by Tukey’s post-hoc test, significance is indicated by orange (5-mm versus 12-mm), blue (5-mm versus 25-mm), and green (12-mm versus 25mm) asterisks. See Supplementary Tables S10-S20 for complete statistical and PCA results.

3-month-old mice significantly changed their locomotor program to adapt to narrowing beams. When mice crossed the 5-mm beam, the stride and the swing duration shortened (Fig. 4d). Further during swing, the ankle and knee angles showed reduced flexion and increased extension (Fig. 4e,f), suggesting that mice walked with a predominantly extended limb posture, indicative of an adaptive strategy to improve balance on a narrow surface. Importantly, hip kinematics were unchanged (Fig. 4g), indicating that the requirement for increased precision of paw placement is implemented mainly through a finer regulation of the distal joints. The challenge to balance on the narrow beam caused the ankle/knee inversion, typically used to generate forward thrust, to be reduced (Supplementary Fig. 4a-c), necessitating a compensatory increase in joint velocities to generate sufficient propulsive strength (Supplementary Fig. 4d-f). Coalescing all meaningful kinematic features (forelimb and hindlimb height, angles and velocity) in PCA space revealed segregated clusters corresponding to the different beam widths (Fig. 4h). Thus, AutoGaitA’s multi-level analysis revealed the kinematic adaptation of locomotor programs to increased task difficulty.

Next, we investigated how age affects these locomotor adaptation strategies by comparing how 3- and 8-month-old mice performed on the beam task. Aging did not change the kinematic pattern induced by the narrow beam, with older mice still showing reduced knee/ankle inversion (Supplementary Fig. 4g-i), increased joint velocity (Supplementary Fig. 4j-l), shortened stride (Supplementary Fig. 4m), less flexed ankle (Fig. 4i) and knee (Supplementary Fig. 4n), and no changes to the hip angle (Supplementary Fig. 4n). However, 8-month-old mice adopted an even more extended limb posture, with the ankle having higher peaks of extension at the end of swing (Fig. 4i). Thus, 8-month-old mice preserve the ability to execute the 25-, 12-, and 5-mm locomotor programs, but with minor adjustments, such as increased distal joint extension (Fig. 4i, Supplementary Fig. 4g-n). Consistently, the PCA showed that the 25-, 12-, and 5-mm locomotor programs cluster separately, but with almost no difference between ages (Supplementary Fig. 4o).

Finally, we compared 8- and 24-month-old mice on the beam task. Age reduced paw placement dexterity, with older mice displaying increased footslips (Supplementary Fig. 5a). Nevertheless, the kinematic adaptation to narrower beams was preserved, as older mice still exhibited shortened stride (Fig. 4j), extended leg posture (Fig. 4k, Supplementary Fig. 5d-f), and increased joint velocity (Supplementary Fig. 5g-i). In contrast to younger mice, 24-month-old mice displayed increased hip flexion on the 5-mm beam (Supplementary Fig. 5c), suggesting a more crouched posture to lower the centre of mass and improve postural stability. This crouched posture was also evident when comparing 8- and 24-month-old mice, as older mice exhibited a reduced peak of ankle extension during swing (Fig. 4k). Finally, the PCA showed a clear separation by both age and beam width (Fig. 4l), suggesting that locomotor programs undergo a more significant change between 8 and 24 months than at earlier stages (Supplementary Fig. 4o).

In summary, AutoGaitA enabled us to identify task-specific locomotor programs, and assess how they change as a function of age.

## Discussion

We have developed AutoGaitA, the open-source Python toolbox providing a versatile framework for the quantitative and standardised assessment and comparison of motor programs at different levels of granularity across species and behaviours. Showcasing the potential of this framework, we used it to reveal three key principles in motor control: i) the species-specific biomechanics underlying propulsive strength generation during walking (Fig. 2); ii) the age-dependent adaptation mechanisms to generate propulsive strength (Fig. 3); iii) the combined effects that concomitant perturbations exert on locomotor programs to preserve robustness and flexibility (Fig. 4).

### A versatile quantitative framework for motor control

One of the key factors underlying the deep-learning posture tracking revolution has been the algorithmś applicability to any species or task^4,5^. We have developed a similarly versatile tool that enables comparable and standardised kinematic outputs to be generated independently of the input. This versatility has already been leveraged to identify gait alterations in cerebellar neurodegeneration^18^, assess to which extent physiological locomotor programs are restored in a mouse model of ataxia^19^, and characterise how perturbations of spinal interneurons affect limb movement in neonatal mice^20^.

AutoGaitA can also be used to assess the motor programs underlying other rhythmic behaviours, such as those commonly used as readouts of dexterity (rope pulling, ladder walking) or deregulated sensation (grooming, chronic scratching). Compiling kinematic data of all these behaviours with AutoGaitA will establish a catalogue of motor programs across model organisms. Moreover, the annotation tables provide an additional tool to compare successful to failed cycles or failed cycles across mutations or perturbations, granting important insight into how motor programs are adapted or degraded by physiological and pathological states. The standardised comparison of motor programs in health and disease will allow to better classify motor disorders, to identify motor symptom progression over the course of diseases and to unveil early disease hallmarks in pre-symptomatic stages.

Beyond, future applications of AutoGaitA could involve assessing athletes’ performance in sports science, optimising treatments in physiotherapy and neurology, studying the impact of affective disorders on motor behaviours in psychiatry, and establishing kinematic readouts of environmental influences (e.g. stress/anxiety) in behavioural research.

### Propulsion in locomotion: species-, age- and task-dependent mechanisms

The generation of propulsion during locomotion varies widely across species, reflecting differences in biomechanical constraint, body size, and environmental demands^21^. In hexapedal organisms such as fruit flies, posterior leg propulsion relies on the synchronised flexion and extension of all distal leg joints, producing a coordinated power stroke that drives forward movement (Fig. 2g,j, Supplementary Fig. 2a,d). Quadrupedal mammals such as mice use a hip-driven initial power stroke, while the coordinated but opposing ankle and knee movements create the primary thrust (Fig. 2h,k, Supplementary Fig. 2b,e). In contrast, humans combine hip oscillations with ankle push-offs, with the ankle generating additional power during the stance-to-swing transition (Fig. 2i,l, Supplementary Fig. 2c,f). Taken together, our data show how evolution has shaped distinct kinematic solutions to the fundamental challenge of efficient forward propulsion of the leg. Notably, in multi-legged species, interlimb coordination is required to preserve postural stability and synergise the propulsion generated by individual limbs^1^.

Ageing degrades locomotor ability in all species by causing declines in muscle strength, joint flexibility, and neural control^22–24^. Yet, whether the age-dependent changes in locomotor programs follow convergent or divergent mechanisms across species has been a long-standing open question. Difficulties in addressing this question arise from studies focusing on different outputs – strength, stride, coordination, endurance, dexterity – and results needing to be evaluated in light of the distinct biomechanics. By using comparable paradigms and standardised readouts, we determined a conserved mechanism of age-dependent locomotor adaptation in flies, mice and humans: the decline of propulsive strength (Fig. 3). This decline manifested within the constraints of each species’ unique biomechanics (Fig. 3). Defining the mechanistic bases underlying the ageing decline of propulsive strength is particularly important with respect to the motor impairments caused by neurodegenerative diseases and for establishing meaningful comparisons between patients and animal models. For example, ankle push-off power is decreased in ageing humans (Fig. 3f)^25,26^, especially in stroke patients^27^. However, our findings imply that in a mouse model of stroke, the loss of propulsive strength should be studied with a focus on the knee/ankle inversion instead of the ankle push-off.

When locomotor tasks become more demanding, such as crossing narrow paths, the nervous system has to balance stiffness and forward propulsion in a way that guarantees postural stability^28^. As mice cross narrow beams, hindlimbs become rather stiff, with propulsive strength being generated by faster joint movements instead of the typical large angular excursions (Fig. 4a-f, Supplementary Fig. 4a-f). We additionally showed that as mice age and require an even finer balance control, this alternative strategy to generate propulsion was complemented by a lower centre of mass (Fig. 4k, Supplementary Fig. 5).

In summary, AutoGaitA represents a much-needed standardized framework for the quantitative assessment of motor programs across behaviours, perturbations, and species. Through our showcase studies, we demonstrated that AutoGaitA enables the discovery of fundamental principles governing motor control across species. We anticipate that this framework will have profound implications for advancing therapeutic interventions, optimizing rehabilitative strategies, and elucidating the evolutionary basis of motor control.

## Acknowledgements

We thank Nicholas Del Grosso (iBots, University of Bonn) for help with research software engineering, Prof. Kathrin Möllenhoff (University of Cologne) for consulting on the appropriate statistical analysis, and the student assistants Sarah Sabbagh and Luca Flemming for their contributions to the code. This work was funded by the Deutsche Forschungsgemeinschaft (DFG, German Research Foundation) - SFB 1451 Project-ID 431549029-INF, 431549029-Z02, 431549029-Z03. M.Ha., A.B., I.K. and G.G. are members of the “iBehave” network funded by the Ministry of Culture and Science of the State of North Rhine-Westphalia.

## Author contribution

Conceptualisation: M.H., I.K., A.G., S.D. and G.G.; data curation: M.H., I.K., V.W., C.S., T.D.K., M.Ha.; analysis: M.H., I.K., V.W., A.K., M.T., M.Ha.; investigation: M.H., I.K., V.W., M.Ha., C.S., T.D.K., V.M., A.G.; methodology: M.H., I.K., V.W., M.Ha., S.D., G.G.; resources: T.K., A.B., P.H.W., C.G., G.R.F., S.D., G.G.; software: M.H.; supervision: T.K., A.B., P.H.W., G.R.F., S.D., G.G.; validation: M.H., I.K., V.W., M.Ha., M.T., G.G.; visualisation: M.H., G.G.; writing of the original draft: M.H., G.G.; reviewing and editing: M.H., I.K., V.W., C.S., T.D.K., V.M., M.Ha., M.T., T.K., A.B., C.G., P.H.W., G.R.F., S.D., G.G.

## Methods

### Software Availability

AutoGaitA is provided as an open-source Python toolbox (GitHub – mahan-hosseini/AutoGaitA: Automated Gait Analysis in Python), being developed on top of well established, documented, and maintained Python dependencies: NumPy^29,30^ (https://numpy.org), SciPy^31^ (https://scipy.org), pandas^32^ (https://pandas.pydata.org), Scikit-Learn^33^ (https://scikit-learn.org), Pingouin^34^ (https://pingouin-stats.org/), Matplotlib^35^ (https://matplotlib.org), seaborn^36^ (https://seaborn.pydata.org), CustomTkinter (https://customtkinter.tomschimansky.com), Pillow^37^ (https://python-pillow.github.io), openpyxl (https://openpyxl.readthedocs.io/en/stable/), ffmpeg-python (https://kkroening.github.io/ffmpeg-python/), and h5py^38^ (https://www.h5py.org).

AutoGaitA’s source code can be accessed in the GitHub repository under the GPLv3 license, and is further integrated with Zenodo^39^ (http://doi.org/10.5281/zenodo.15373063), automatically linking all releases to unique digital object identifiers (DOIs). AutoGaitA’s central goal is to take off the burden of post-tracking analyses from researchers, particularly those not too familiar with programming. We thus provide easy-to-understand graphical user interfaces (GUIs), <u>a</u> <u>straightforward documentation</u> consisting of many images, and comprehensive video tutorials on the <u>AutoGaitA YouTube channel</u>. Nonetheless, we have simultaneously made it easy for well-versed developers and users to contribute to and extend AutoGaitA as well as to integrate our toolbox into custom workflows and other (Python) tools. We strongly appreciate and encourage user feedback and contributions via or directly as GitHub pull requests.

### AutoGaitA workflow details

#### Data preparation and important settings

For AutoGaitA DLC and SLEAP, we provide two ways of naming input data files: a thorough and safe way, which provides detailed information about the contents of files, and a reduced and risky way, which is quick to implement but carries some ambiguity. AutoGaitA Universal 3D includes a tool which allows convenient adaptation of the required naming convention of the columns of corresponding 3D data files. Since AutoGaitA Universal 3D analyses 2D kinematics of 3D body coordinates, users are advised to pay attention to their behaviour of interest and to which dimension’s features are of particular interest (e.g. along which 2D-plane angles should be computed). Excel tables containing the timestamp annotations of the behaviour to be analysed (annotation tables) need to be prepared before any analysis. These need to follow a specific format, which differs slightly between AutoGaitA DLC/SLEAP and AutoGaitA Universal 3D.

#### Normalisation of segmented data

In first-level workflows (Figure 1b), segmented data is normalised to be equally long for all cycles. In the explanations below, we will use *time-points* when referring to the length of original cycles before normalisation and *cycle-bins* when referring to the length of normalised cycles. Cycles are normalised to a user-chosen target length n (in cycle-bins) by being either extended if they were shorter (i.e. had fewer time-points than n originally) or compressed if they were longer (i.e. had more time-points than n originally). For example, if the user selected to normalise to ten cycle-bins, all original cycles of length 9 or below would be extended, and all original cycles of length 11 or more would be compressed. Extension is done via repetition. For the example target value of 10 cycle-bins and an original cycle having only 5 time-points before normalisation, all its data values will be repeated once (i.e.: original data at time-points: 1, 1, 2, 2, … 5, 5 = normalised data at cycle-bins 1, 2, 3, 4, … 9, 10). Compression is done via averaging. In our example, if the original cycle comprises 20 time-points, its data points are averaged in adjacent pairs: first and second, third and fourth data points and so on (i.e.: averages of data at time-points: 1 & 2, 3 & 4, 5 & 6, 7 & 8, … 17 & 18, 19 & 20 = normalised data at cycle-bins 1, 2, 3, …, 9, 10).

After normalisation, each cycle-bin thus represents the data at respective percentiles, based on the number of selected bins. Default settings normalise to 25 cycle-bins, leading to cycle-bins reflecting 4% percentiles: 1-4%, 5-8%, 9-12%, etc., since this value showed the best comparability in our cross-species locomotion analysis.

#### Standardisation of direction of movement

In the same locomotion experiment, animals might walk through the video screen from left to right or vice versa. To ensure that the data is independent of the direction of motion, AutoGaitA offers options to adjust the horizontal coordinates (e.g., x-coordinates in 2D), effectively simulating that all animals walk towards the same point. For behaviours like jumping jacks or rope climbing, which are approximately stationary along the horizontal dimension, this option should be turned off. Users of AutoGaitA Universal 3D should consider additional details about this standardisation in 3D in the documentation’s section on *back and forth behaviours*.

#### Standardisation of horizontal coordinates

AutoGaitA provides an option to analyse horizontal coordinates, for example, x-coordinates in 2D (see Fig. 4d). In a locomotion study, the x-coordinates of the foot inform about its distance travelled with each step (i.e., the stride). To ensure that x-coordinates are comparable across steps, they must be standardised to the same position across cycles. This standardisation ensures that no bias is introduced by AutoGaitA averaging features across cycles (and, consecutively, across animals at the group level). For standardisation, we subtract, at each cycle separately, the minimum x-coordinate of a user-chosen key point from those of all key points. As a result, x-coordinates after standardisation inform about key points’ distance to the standardisation-joint’s zero-point and are, critically, independent of their absolute position in space.

#### Standardisation of vertical coordinates

Vertical coordinates, for example y-coordinates in 2D, can be standardised according to baseline, global, or landmark standardisation. In the (typically most accurate) baseline standardisation, the y-coordinates of a tracked reference baseline, like the beam in our mouse experiments (Fig. 2b), are subtracted from those of the animal’s body at the corresponding time points. If no baseline data is available, users can use global or landmark standardisation. Global standardisation subtracts the smallest y-coordinate present across the full dataset (i.e., across time and all landmarks) from the y-coordinates of all body landmarks. Landmark standardisation subtracts the smallest y-coordinate of a user-chosen landmark from the body’s y-coordinates. AutoGaitA also provides an option to standardise the y-coordinates of each step separately, which is recommended if no baseline standardisation is possible and the floor is uneven or cameras are distorted.

#### Principal Components Analysis

Users can choose which kinematic features to include in AutoGaitA’s Principal Components Analysis (PCA), which commences with extracting averages of these features across the entire cycle for each animal (Fig. 1c). Each cycle-bin of each feature is included as an input feature in the PCA. For example, if the ankle angle is an input feature and cycles were normalised to 10 cycle-bins, the PCA would have 10 input features capturing animals’ ankle angles: ankle angle 1-10% cycle, ankle angle 11-20% cycle, …, ankle angle 91-100% cycle. Users have the option to compute input features only over a subset of the cycle (e.g. the first half, or the first and last quartile) instead. Following the PCA convention, input features are standardised to have zero mean and unit variance before the model is fitted. Scikit-Learn^33^ is used for standardisation as well as model-fitting. Depending on user input, the number of returned principal components (PC) can either be chosen directly or configured to explain a certain percentage of data variance. AutoGaitA produce5s PCA outputs as: 2D- and 3D scatterplots, generating video-files of the latter if wanted, and tabular files containing PC’s explained variance, input features’ eigenvectors, each animal’s coordinates in PC-space, and an overview of the 20 input features that contributed most strongly to each PC to simplify their interpretability.

#### One-way ANOVA and Tukey’s Test

AutoGaitA provides between- or within-subjects one-way ANOVAs, with the former assessing different subjects, as in Fig. 3, and the latter assessing the same subjects across different conditions, as in Fig. 4d-f, for example. ANOVA results are provided as conventional ANOVA tables in text files. While the one-way ANOVA tests a certain feature for group differences globally, Tukey’s post-hoc test compares the feature at individual cycle-bins separately, correcting for the number of multiple comparisons (AutoGaitA does not require the ANOVA for Tukey’s to be run). Besides figures illustrating the results of Tukey’s tests, we provide their exact numerical results (i.e., Tukey’s q-values as well as corresponding p-values and confidence intervals) in text and tabular files. AutoGaitA uses Pingouin^34^ for one-way ANOVAs and SciPy^31^ for Tukey’s tests.

#### Cluster-extent Permutation Test

The cluster-extent test is preferred over the ANOVA whenever parametric assumptions are not met. AutoGaitA’s cluster-extent test follows the concepts introduced by Maris and Oostenveld^40^, which are by now well-established in the field of human electrophysiology. We provide an in-depth explanation of how the test is implemented in AutoGaitA in Hosseini et al.^41^. The outputs of the cluster-extent test are provided as figures as well as text files, storing the p-values of all clusters. AutoGaitA uses SciPy^31^ for t-tests and Scikit-Learn^33^ for shuffling data randomly.

### Experimental fly model

Adult male wild-type Canton-S *Drosophila melanogaster* flies were collected after eclosion and reared on a standard yeast-based medium^42^ at 25°C and 65% humidity in a 12-hour dark/light cycle. Experiments were performed with young (2-3 days old) and old (21-22 days old) flies. Tethered flies walked on a spherical treadmill and were recorded from the side using a high-speed camera (acA1300-200um, Basler), equipped with a 50 mm lens (LM50JC1MS, Kowa Optical Products)^12^. Videos were recorded at 400 frames/second with a resolution of 912 x 550 pixels. To convert pixels into metric scale, the camera was calibrated with a custom-made checkerboard pattern (7 x 6 squares with size 399 µm x 399 µm per square) developed on a photographic slide. The conversion factor of 5.883 ± 0.198 µm per pixel (mean ± standard deviation) was determined by analysing the side length of 1850 squares from 74 images of the checkerboard.

DeepLabCut^4^ was used for automated tracking of leg landmarks in the videos by training a ResNet-50 network with a training set containing 755 images (10 videos with 54 to 105 frames each) from 5 flies. Resulting leg landmark predictions were visually inspected and manually corrected as needed with a custom-made graphical user interface software. Begin and end times for swing and stance phases were manually annotated for each step.

### Experimental mouse model

Mice were maintained following the protocols for animal experiments approved by the local health authority in North Rhine-Westphalia (LAVE, Landesamt für Verbraucherschutz und Ernährung, Nordrhein-Westfalen). 3-month-, 8-month- and 24-month-old C56BL6/J mice of both sexes were used for behavioural experiments. These mouse age groups are commonly used to characterise gait following physiological, pathological and circuit perturbations. Analysis of the behavioural data showed similar responses in male and female mice. Mice had *ad libitum* access to food and water, and were housed in groups of maximum 5 animals, maintained on a 12-hour dark/light cycle within a room controlled for humidity and temperature.

We opted for the beam paradigm to test a challenge, e.g. walking on a narrowing path, mice are likely to encounter also in their naturalistic settings. Naïve mice of different ages were tested as they crossed 1.3-meter-long beams with different widths (5-mm, 12-mm, and 25-mm). Studying untrained mice enables the identification of innate adaptation strategies in response to the width-dependent perturbations. On the first day, mice were tasked to cross the wide, 25-mm, beam, on the second day the 12-mm beam, and on the third day the narrow, 5-mm, beam. For each beam size, three to five trials per mouse were recorded using eight high-speed cameras (mV Blue Cougar XD) positioned around the beam (3D Simi Motion). Multiple camera views were analysed to count the number of slips, which were then averaged across trials per individual mouse. Videos were recorded at 100 frames/second, with a resolution of 1200 x 900 pixels.

The Simi™ Motion software (version 10.2.0) was used to record markerless mice walking in both directions, so our data are averages of both right and left limbs. We used DeepLabCut^4^ to track the body landmark coordinates, as well as the beam surface to define our vertical baseline. Tracking was done after training a ResNet-50 network on 2147 frames (113 videos with 19 frames each) from 8 mice. The number of slips and the phases of the step cycle (beginning of swing, end of swing, and end of stance) were manually annotated. Slips or pauses on the beam were excluded from the kinematic analysis. Please note that 24-month-old mice display an increased number of slips on the narrowest beam (5-mm), thus, we analysed fewer animals compared to the other tasks.

### Human dataset

Participants meeting the following criteria were eligible for inclusion in the study: Age between 21 and 90 years, written informed consent, absence of neurological or psychiatric diseases and no health conditions affecting the locomotor system. The study was conducted according to the principles of the Declaration of Helsinki and approved by the local ethics committee of the Faculty of Medicine at University of Cologne (21-1418_1). The final sample is presented in Supplementary Table S32 and consisted of 47 individuals, including 29 younger participants (age [mean ± SD]: 28.1 ± 3.5 years, age range: 21 to 36 years; 19 females) and 18 older participants (age [mean ± SD]: 67.7 ± 11.0 years, age range: 46 to 85; 9 females). There were no significant differences between the two age groups with regard to sex (χ² = 0.560, p = 0.454) and body height (t= 0.819, p = 0.417).

Participants walked on a walkway (460 cm in length x 60 cm in width), while being recorded by eight high-speed cameras (mvBlueCougar XD, Matrix Vision GmbH) positioned in a circular arrangement around the walkway. Videos were recorded at 100 frames/second, with a resolution of 1936 x 1216 pixels. The Simi™ Motion software (version 10.2.0) was used for the video recordings and DeepLabCut^4^ was used to track body landmarks. Tracking was done after training a ResNet-50 network on 4797 frames (123 videos with 39 frames each) from 31 humans.

At the beginning of the recordings, participants were asked to stand at a marked position on one end of the walkway. After a verbal signal, participants started walking at a convenient, self-generated speed to the other end of the walkway. Once they arrived at the end, they turned around and walked back to the start position. This back-and-forth walking was repeated until up to three trials were performed by each participant. Videos were split based on turns before being tracked with DLC, meaning that (as for our mouse data) the data corresponds to averages of left and right legs. Step cycle phases, i.e., the start of the swing phase, the end of the swing phase, and the end of the stance phase, were annotated manually. More specifically, the toe-off moment of the feet was marked as the end of the stance phase and simultaneously as the start of the swing phase, while the moment of the heel-strike was marked as the end of the swing phase and simultaneously as the start of the stance phase.

## References

1. Büschges, A. & Ache, J. M. Motor control on the move: from insights in insects to general mechanisms. Physiol. Rev. 105, 975–1031 (2025).

2. Grillner, S. & El Manira, A. Current Principles of Motor Control, with Special Reference to Vertebrate Locomotion. Physiol. Rev. 100, 271–320 (2020).

3. Bernstein, N. The Co-Ordination and Regulation of Movements. (Pergamon Press; Oxford, 1967).

4. Mathis, A. et al. DeepLabCut: markerless pose estimation of user-defined body parts with deep learning. Nat. Neurosci. 21, 1281–1289 (2018).

5. Pereira, T. D. et al. SLEAP: A deep learning system for multi-animal pose tracking. Nat. Methods 19, 486–495 (2022).

6. Aljovic, A. et al. A deep learning-based toolbox for Automated Limb Motion Analysis (ALMA) in murine models of neurological disorders. Commun. Biol. 5, 131 (2022).

7. Lv, X. et al. PMotion: an advanced markerless pose estimation approach based on novel deep learning framework used to reveal neurobehavior. J. Neural Eng. 20, 046002 (2023).

8. Tozzi, F., Zhang, Y.-P., Narayanan, R., Roquiero, D. & O’Connor, E. C. Forestwalk: A machine learning workflow brings new insights into posture and balance in rodent beam walking. bioRxiv 2024.04.26.590945 (2024) doi:10.1101/2024.04.26.590945.

9. Ruiz-Vitte, A. et al. Ledged Beam Walking Test Automatic Tracker: Artificial intelligence-based functional evaluation in a stroke model. Comput. Biol. Med. 186, 109689 (2025).

10. Chockley, A. S. et al. Subsets of leg proprioceptors influence leg kinematics but not interleg coordination in *Drosophila melanogaster* walking. J. Exp. Biol. 225, jeb244245 (2022).

11. DeAngelis, B. D., Zavatone-Veth, J. A. & Clark, D. A. The manifold structure of limb coordination in walking Drosophila. eLife 8, e46409 (2019).

12. Haustein, M., Blanke, A., Bockemühl, T. & Büschges, A. A leg model based on anatomical landmarks to study 3D joint kinematics of walking in Drosophila melanogaster. Front. Bioeng. Biotechnol. 12, 1357598 (2024).

13. Pratt, B. G., Lee, S.-Y. J., Chou, G. M. & Tuthill, J. C. Miniature linear and split-belt treadmills reveal mechanisms of adaptive motor control in walking Drosophila. Curr. Biol. 34, 4368–4381.e5 (2024).

14. Bellardita, C. & Kiehn, O. Phenotypic Characterization of Speed-Associated Gait Changes in Mice Reveals Modular Organization of Locomotor Networks. Curr. Biol. 25, 1426–1436 (2015).

15. Takeoka, A., Vollenweider, I., Courtine, G. & Arber, S. Muscle Spindle Feedback Directs Locomotor Recovery and Circuit Reorganization after Spinal Cord Injury. Cell 159, 1626–1639 (2014).

16. Gonzalez-Islas, J. C., Dominguez-Ramirez, O. A., Castillejos-Fernandez, H. & Castro-Espinoza, F. A. Human gait analysis based on automatic recognition: A review. Pädi Bol. Científico Cienc. Básicas E Ing. ICBI 10, 13–21 (2022).

17. Ghorbani, S. et al. MoVi: A large multi-purpose human motion and video dataset. PLOS ONE 16, e0253157 (2021).

18. Tolve, M. et al. The endocytic adaptor AP-2 maintains Purkinje cell function by balancing cerebellar parallel and climbing fiber synapses. Cell Rep. 44, (2025).

19. Tutas, J. et al. Autophagy regulator ATG5 preserves cerebellar function by safeguarding its glycolytic activity. Nat. Metab. 7, 297–320 (2025).

20. Trevisan, A. J. et al. The transcriptomic landscape of spinal V1 interneurons reveals a role for En1 in specific elements of motor output. Preprint at 10.1101/2024.09.18.613279 (2024).

21. Schaeffer, P. J. & Lindstedt, S. L. How Animals Move: Comparative Lessons on Animal Locomotion. in Comprehensive Physiology (ed. Prakash, Y. S.) 289–314 (Wiley, 2013). doi:10.1002/cphy.c110059.

22. Desforges, J. F. & Sudarsky, L. Gait Disorders in the Elderly. N. Engl. J. Med. 322, 1441–1446 (1990).

23. Ingram, D. K., London, E. D., Reynolds, M. A., Waller, S. B. & Goodrick, C. L. Differential effects of age on motor performance in two mouse strains. Neurobiol. Aging 2, 221–227 (1981).

24. Jones, M. A. & Grotewiel, M. Drosophila as a model for age-related impairment in locomotor and other behaviors. Exp. Gerontol. 46, 320–325 (2011).

25. Franz, J. R. The Age-Associated Reduction in Propulsive Power Generation in Walking. Exerc. Sport Sci. Rev. 44, 129–136 (2016).

26. Sloot, L. H. et al. Decline in gait propulsion in older adults over age decades. Gait Posture 90, 475–482 (2021).

27. Balasubramanian, C. K., Bowden, M. G., Neptune, R. R. & Kautz, S. A. Relationship Between Step Length Asymmetry and Walking Performance in Subjects With Chronic Hemiparesis. Arch. Phys. Med. Rehabil. 88, 43–49 (2007).

28. Earhart, G. M. Dynamic control of posture across locomotor tasks. Mov. Disord. 28, 1501–1508 (2013).

29. Harris, C. R. et al. Array programming with NumPy. Nature 585, 357–362 (2020).

30. Van Der Walt, S., Colbert, S. C. & Varoquaux, G. The NumPy Array: A Structure for Efficient Numerical Computation. Comput. Sci. Eng. 13, 22–30 (2011).

31. Virtanen, P. et al. SciPy 1.0: fundamental algorithms for scientific computing in Python. Nat. Methods 17, 261–272 (2020).

32. McKinney, W. Data Structures for Statistical Computing in Python. in 56–61 (Austin, Texas, 2010). doi:10.25080/Majora-92bf1922-00a.

33. Pedregosa, F. et al. Scikit-learn: Machine Learning in Python. J Mach Learn Res 12, 2825–2830 (2011).

34. Vallat, R. Pingouin: statistics in Python. J. Open Source Softw. 3, 1026 (2018).

35. Hunter, J. D. Matplotlib: A 2D Graphics Environment. Comput. Sci. Eng. 9, 90–95 (2007).

36. Waskom, M. seaborn: statistical data visualization. J. Open Source Softw. 6, 3021 (2021).

37. Murray, Andrew et al. python-pillow/Pillow: 11.2.1. Zenodo 10.5281/ZENODO.596518 (2025).

38. Collette, A. Python and HDF5: Unlocking Scientific Data. (O’Reilly, Beijing Köln, 2014).

39. Hosseini, M. AutoGaitA - Automated Gait Analysis in Python. Zenodo 10.5281/ZENODO.15373063 (2025).

40. Maris, E. & Oostenveld, R. Nonparametric statistical testing of EEG- and MEG-data. J. Neurosci. Methods 164, 177–190 (2007).

41. Hosseini, M. et al. AutoGaitA – Automated Gait Analysis in Python. bioRxiv 2024.04.14.589409 (2024) doi:10.1101/2024.04.14.589409.

42. Backhaus, B.S.E., Sulkowski, E. & Schlote, F.W. A semi-synthetic, general-purpose medium for Drosophila melanogaster. in Drosophila Information Service vol. 60 210–212 (1984).

