## Supplementary Note 1 for "AutoGaitA: A versatile quantitative framework for kinematic analyses across species, perturbations and behaviours"

### Supplementary Note 1: Adopting AutoGaitA across behaviours with the MoVi dataset

In this section we will adopt the AutoGaitA workflow (Fig. 1) on the open-source MoVi dataset[1] to demonstrate that our toolbox is effective in identifying the motor strategies of a wide range of behaviours, not just locomotion. This analysis further illustrates that AutoGaitA Universal 3D can be used with any kind of (marker-based or marker-less) 3D coordinate data, given that coordinates are provided in Excel tables.

To sum, MoVi videos and MATLAB-datasets are provided as a single sequence of 21 behaviours. The dataset's authors provide scripts to split these sequences into separate behaviours (see https://github.com/saeed1262/MoVi-Toolbox). Based on these scripts we wrote custom MATLAB scripts that generate coordinate Excel files for each subject and each behaviour. The following behaviours were included: *walking, running, crawling, sideways (walking), vertical jumping, and jumping jacks*. This selection reflects the fact that AutoGaitA workflow is particularly suited for rhythmic behaviours, i.e., behaviours with similar temporal characteristics across repetitions and individuals. Non-rhythmic behaviours provided with MoVi were, for example, *dj-ing* or *taking photos*. AutoGaitA Universal 3D's *Datafile Preparation* (see our documentation for details) was then used to ensure that the columns of Excel files corresponded to the naming convention required by AutoGaitA. Next, annotation tables of cycles were manually generated as usual.

AutoGaitA Universal 3D's configuration was adjusted to reflect MoVi's settings (e.g., a sampling rate of 120 Hz) and were fixed across behaviours with one exception. The setting to simulate equal gait direction was turned on for all behaviours except vertical jumping and jumping jacks, since these behaviours are approximately stationary on the y-dimension. In the MoVi dataset, coordinates that could not be tracked are assigned a zero. To handle conflicts with coordinates really being zero, we added a check that excluded cycles from AutoGaitA

Universal 3D's analysis if at least two consecutive time points had a value of zero (which is highly unlikely for real coordinates but was likely always the case for tracking failures). This excluded the crawling behaviour, since only two usable cycles were left across all subjects for it. The final number of subjects of the other five behaviours were: 82 for running & sideways, 85 for walking, 87 for vertical jumping & jumping jack.

The PCA was configured to fit 10 components on the y- and z-coordinates of the head, hip, pelvis, thorax, ankles, elbows, hips, hands, knees, shoulders, wrists, and foots. Angles of the ankles, knees, elbows, and wrists, were included as well as all corresponding velocities and accelerations. PCA results are provided in Supplementary Fig. 1, with panels a, b, and c illustrating the distribution of explained variance per component as well as scatterplots in two- and three-dimensions, respectively. Supplementary Fig. 1b,c demonstrate clear clustering of behaviours based on their motor signatures with behaviours that are more similar (e.g., walking and running) being closer in Principal Component (PC)-space, too. This is in contrast to sideways walking, which is clearly most distinct to the other four behaviours examined and, as a result, has a distinct cluster in PC-space. It is further striking that the most similar behaviours of vertical jumping and jumping jacks, despite clustering very closely in PC-space, barely overlap. This pattern nicely signifies that even subtle differences between motor signatures are identified well.

### References

1. Ghorbani, S. *et al.* MoVi: A large multi-purpose human motion and video dataset. *PLOS ONE* **16**, e0253157 (2021).

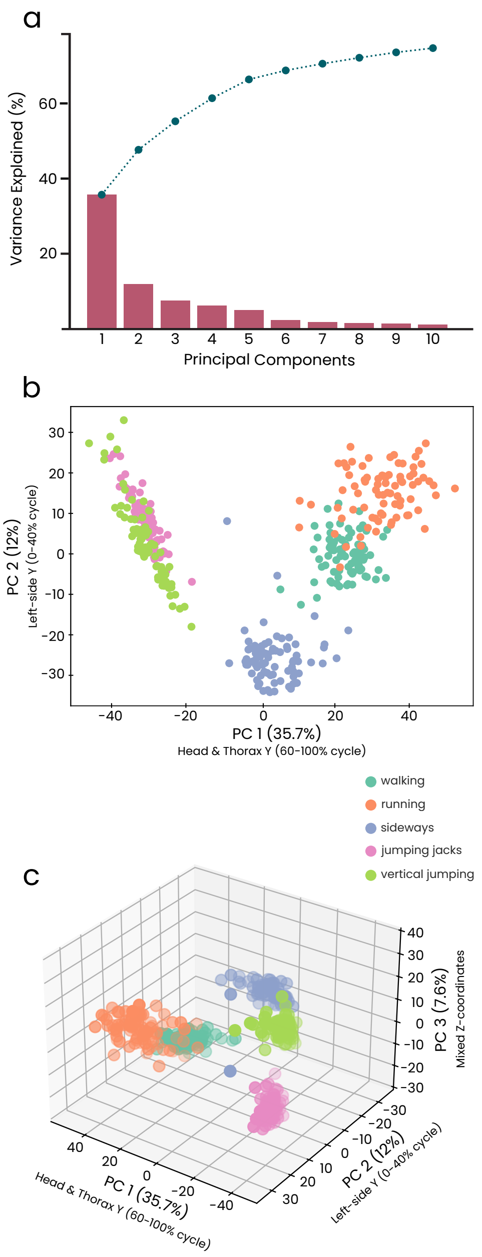


**Supplementary Fig. 1 | AutoGaitA's PCA clusters the motor program underlying different human rhythmic behaviours in segregated spaces.**

**a,** The absolute and cumulative proportion of variance explained by each of the ten fitted principal components (PCs). **b,c,** 2D (b) and 3D (c) PCA scatterplots showing the segregation of 5 distinct rhythmic behaviours, with each cluster representing a motor program. N=82 subjects, each depicted as a coloured circle. Scatterplots show that behaviours are successfully clustered based on PC 1 (36% variance explained), reflecting Head/Thorax-Y coordinates in the second half of the cycle (60-100%), PC 2 (12% variance explained), reflecting early (0-40% cycle) left-side Y-coordinates, and PC 3 (7.6% variance explained), reflecting mixed Z-coordinates. Complete PCA results are provided in Supplementary Tables S21 & S22.

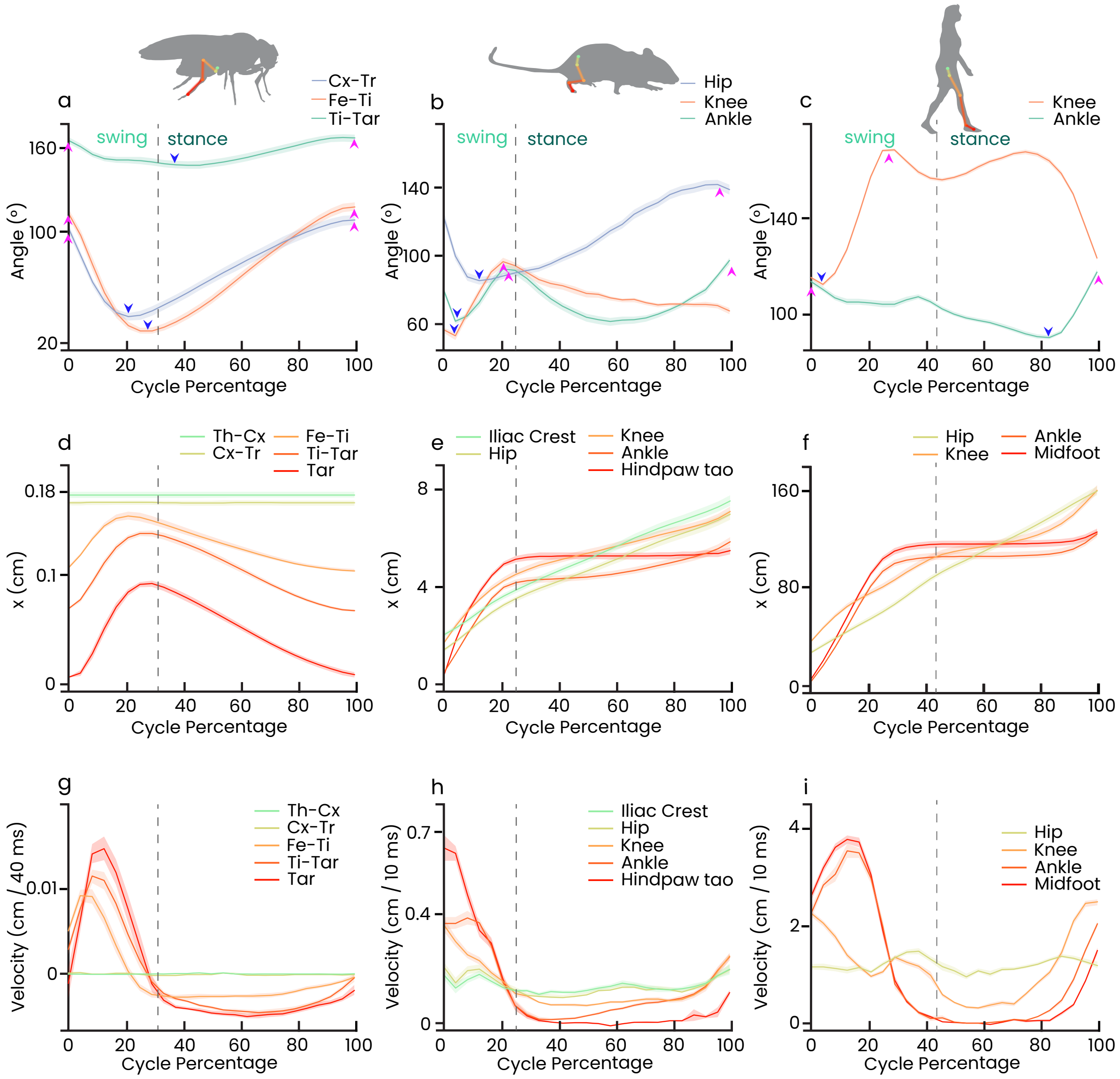


**Supplementary Fig. 2 | Kinematic assessments of the locomotor programs of the lower limbs across species.**
**a,** Excursions of the coxa-trochanter (Cx-Tr), femur-tibia (Fe-Ti), tibia-tarsus (Ti-Tar) angles throughout the step cycle of the hind leg in flies. Peak extension and flexion of the angles in a-c are indicated with purple and blue arrowheads, respectively. **b,** Excursions of the ankle, knee, and hip angles throughout the step cycle in mice. **c,** Excursions of the ankle and knee hip angles throughout the step cycle in humans. **d-f,** Variations in X-coordinates of selected landmarks on the fly hind leg (d), the mouse hindlimb (e) and the human leg (f). **g-i,** Horizontal velocities (i.e., along x-dimension) of fly (g), mouse (h) and human (i) landmarks. N=6 flies, N=9 mice and N=29 human subjects. Data are presented as mean±SEM, SEM is represented as shaded areas.

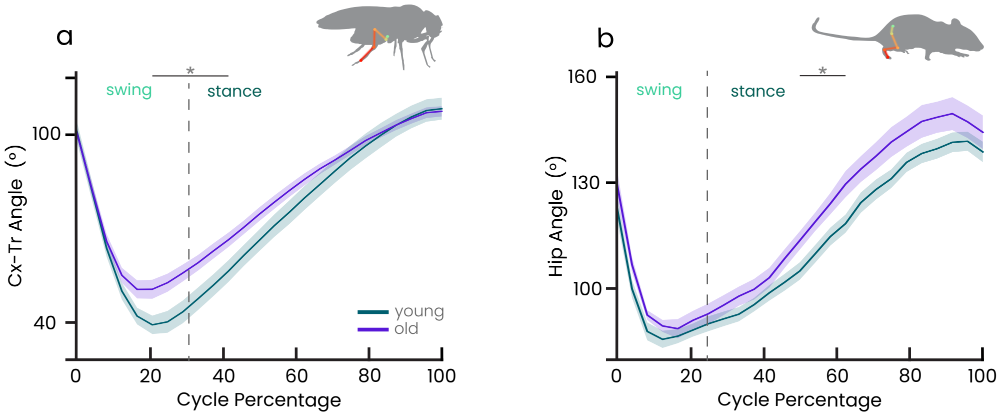


**Supplementary Fig. 3 | Age-dependent effects on angles in flies and mice.**
**a,** The coxa-trochanter (Cx-Tr) angle in older flies is less flexed at the swing-stance transition. N=6 young (green, 2-3 days) and N=6 old (purple, 21-22 days) flies. **b,** Hip angle in older mice is slightly less flexed during mid-stance. N=9 young (green, 8 months), and N=12 old (purple, 24 months) mice. Data are presented as mean±SEM, SEM is represented as shaded areas. Statistical analysis: one-way ANOVA followed by Tukey's post-hoc, significance is indicated with asterisks, * $p<0.05$. See Supplementary Tables S23 & S24 for complete statistical results.

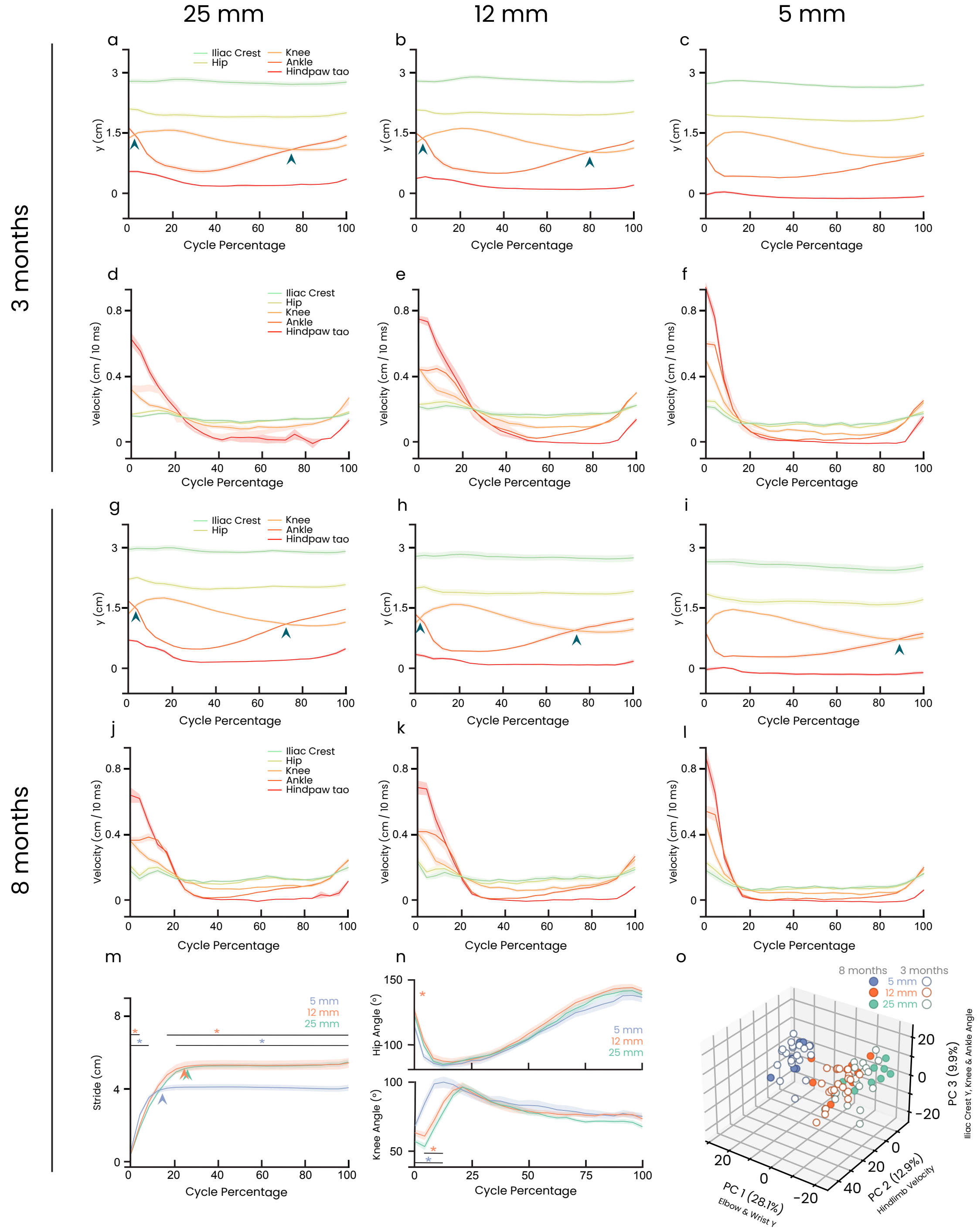


**Supplementary Fig. 4 | Kinematic changes in 3- and 8-month-old mice walking on 25 mm, 12 mm and 5 mm wide beams.** **a-c,** Changes in hindlimb landmark height through the step cycle, as 3-month-old mice cross a 25 mm (a), 12 mm (b) and 5 mm (c) wide beam. Dark cyan arrowheads indicate ankle / knee inversion. **d-f,** Changes in hindlimb landmark velocity through the step cycle, as 3-month-old mice cross a 25 mm (d), 12 mm (e) and 5 mm (f) wide beam. **g-i,** Changes in hindlimb landmark height through the step cycle, as 8-month-old mice cross a 25 mm (g), 12 mm (h) and 5 mm (i) wide beam. **j-l,** Changes in hindlimb landmark velocity through the step cycle, as 8-month-old mice cross a 25 mm (j), 12 mm (k) and 5 mm (l) wide beam. **m,** Changes in stride, measured as distance covered by the hindpaw on the x-axis, through the step cycle, as 8-month-old mice cross a 25 mm (j), 12 mm (k) and 5 mm (l) wide beam. The coloured arrowheads indicate the shortened stride. **n,** Changes in hindlimb knee and hip angles through the step cycle, as 8-month-old mice cross different beams. **o,** PCA of changes in heights, angles and velocities of the forelimb and hindlimb throughout the step cycle between 8-month-old (solid dots) and 3-month-old (framed dots) mice revealed segregated clusters based on beam widths but not age. Data presented as mean±SEM, 3-month-old N=20, 8-month-old N=9. Statistical analysis: one-way ANOVA followed by Tukey's post-hoc, significance is indicated by orange (5-mm versus 12-mm), blue (5-mm versus 25-mm), and green (12-mm versus 25mm) asterisks. See Supplementary Tables S25-S29 for complete statistical and PCA results.

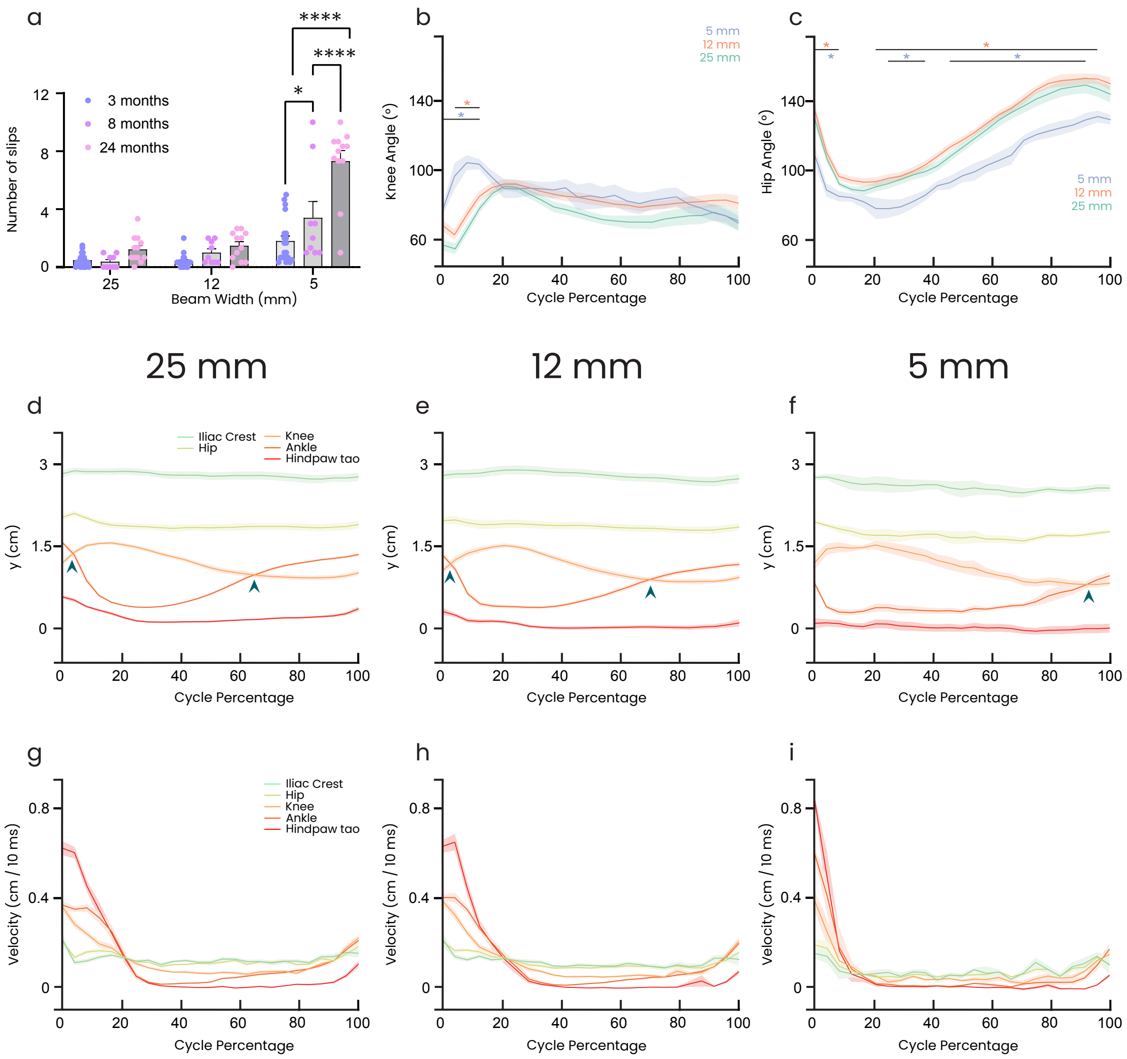


**Supplementary Fig. 5 | Kinematic changes in 24-month-old mice walking on 25 mm, 12 mm and 5 mm wide beams.** **a,** 24-month-old mice show more footslips on the 5 mm beams compared to younger mice. **b,c,** Changes in hindlimb knee (b) and hip (c) angles through the step cycle, as 8-month-old mice cross different beams. **d-f,** Changes in hindlimb landmark height through the step cycle, as 24-month-old mice cross a 25 mm (d), 12 mm (e) and 5 mm (f) wide beam. Dark cyan arrowheads indicate ankle / knee inversion. **g-i,** Changes in hindlimb landmark velocity through the step cycle, as 24-month-old mice cross a 25 mm (g), 12 mm (h) and 5 mm (i) wide beam.Data presented as mean±SEM, 24-month-old N=12. Statistical analysis: one-way ANOVA followed by Tukey's post-hoc, significance is indicated by orange (5-mm & 12-mm), blue (5-mm & 25-mm), and green (12-mm & 25mm) asterisks. See Supplementary Tables S30 & S31 for complete statistical results.
